# Continuous Topological Accessibility (RFA) as a 3D-Aware GRN Filter: Anticorrelation with eQTL and No Prediction of ΔRNA

**DOI:** 10.64898/2026.11.02.763620

**Authors:** Andrey Timofeev, Alexander Bratishchikov, Alexander Anufriev

## Abstract

We formalize *Relative Forest Accessibility* (RFA), a topological accessibility index computed as the solution of (*I* − *γP*)*x* = *v*_0_, where *P* is a Markov transition matrix built from Hi-C contacts and *v*_0_ is a vector of TF peaks (or ATAC peaks). RFA yields a continuous accessibility profile across all genomic bins.

We test RFA on three targets. First, on promoter TF counts (GM12878 chr1, 10 kb, 811 genes), RFA loses to 1D distance (AUC 0.836 vs 0.976); this is an artifact because the target and the 1D distance look at the same window. Second, on eQTL (GTEx v10 LCL, chr1+17+19, 10 kb, 1625 genes), RFA *anticorrelates* with eQTL: *ρ*(RFA, −log_10_ *p*_min_) = −0.09, AUC(RFA) = 0.386 (below chance), AUC( −RFA) = 0.614. High RFA corresponds to fewer eQTL, and dose-response analysis confirms that AUC(RFA) decreases with eQTL strength (mean trend −0.056, 19/20 TFs). A distance-preserving shuffle of the Hi-C matrix reproduces this anticorrelation (ΔAUC vs real: [ −0.074, +0.076] across 5 TFs, all CIs include 0), whereas a degree-preserving rewiring destroys it (ΔAUC ∈ [ −0.261, − 0.003], CIs exclude 0 for 5/5 TFs); the anticorrelation therefore reflects *distance-decay of Hi-C contacts (local 3D accessibility)*, not higher-order topology. The anticorrelation replicates on chr17 (Δ*ρ* = −0.13 for rewired, 0/5 for shuffled) and on chr19 with an inverted control pattern, consistent with local within-diagonal heterogeneity as the carrier on short gene-dense chromosomes. Third, on ΔRNA (MCF10A vs MCF7, 40 kb, ATAC peaks), RFA does not predict ΔRNA (*ρ* = +0.005, permutation *p* = 0.87, bootstrap 95% CI [ −0.040, +0.053], *n* = 1730); the CI includes zero and does not exclude the practical boundary |*ρ*| = 0.05. Within-line sanity checks are also null.

We conclude that RFA anticorrelates with eQTL via local distance-decay and does not predict ΔRNA. The anticorrelation is carried by distance-decay, not by loops or TADs, and may reflect expression buffering: genes with strong eQTL are less dependent on local 3D accessibility. RFA is not a universal predictor but a filter with an inverted sign on eQTL and no signal on ΔRNA.

## 1 Introduction

### 1.1 Problem

Existing 3D-aware TF→gene filters (SSD 2017, Esearch3D 2023, ChIPSP 2026) use either a binary (loop yes/no) or a scalar (direct contact, 1D distance). RFA offers a continuous accessibility profile. The key question is: *where does RFA give signal, in which direction, and what is the mechanism*.

### 1.2 Positioning

We position RFA not as a “new framework” but as “continuous topological accessibility as a resolution-dependent 3D-aware GRN filter”. RFA is not a universal predictor but a filter with a clearly delineated applicability domain and a *signal direction*.

### 1.3 Contributions

1. Formalization of RFA via a Markov random walk (Section 2).
2. **Anticorrelation of RFA with eQTL** (AUC(−RFA) = 0.614, dose-response trend −0.056).
3. **Negative result on** Δ**RNA** (*ρ* = 0.005, MDR= 0.067).
4. **Mechanism:** the eQTL anticorrelation is carried by distance-decay (topology controls, Section 5.3), not by higher-order topology.
5. **Methodological warning:** an automatic sign flip in AUC can mask anticorrelation as “high AUC”; dose-response reveals it.

## 2 Theory

We formalize the computation of a *topological accessibility index* (RFA) that quantifies how accessible a genomic locus is to a transcription factor (TF) given the 3D chromatin architecture measured by Hi-C. The construction proceeds in four stages: from Hi-C contact counts to a weighted graph; from the graph to a Markov transition matrix; from the transition matrix to a continuous accessibility profile; from the profile to per-gene scores and validation statistics. Throughout, the Markov chain is a *mathematical instrument for deconvolving spatial noise and accounting for multi-step paths*, not a kinetic model of TF motion.

### 2.1 Hi-C Graph Construction

Let 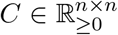 be a symmetric Hi-C contact matrix at resolution *r* (e.g., *r* = 10 kb), where *n* is the number of genomic bins measured by Hi-C [1, 3]. We apply:

1. **Diagonal zeroing**: *C*_*ii*_ := 0 for all *i*, eliminating self-contacts that would otherwise dominate the degree distribution.
2. **Top-***k* **sparsification**: For each row *i*, retain only the *k* largest entries, yielding a sparse matrix *Ã*. This replaces dense noise with the strongest topological signals. Statistical treatment of Hi-C noise and sparsification is discussed in [2].
3. **Symmetrization**: *A* := max(*Ã, Ã*^*⊤*^), preserving symmetry.

A/B compartments (the dominant large-scale chromatin structure) are captured by the first principal component of log(1 + *C*). To isolate fine-scale topological structure from compartment signal, we regress each row of *A* on the compartment score and retain residuals:

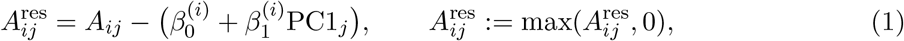

where 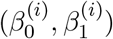 are obtained by ordinary least squares on the support of row *i*.

### 2.2 Markov Transition Matrix

Define the degree vector *d*_*i*_ =∑ _*j*_ *A*_*ij*_ and the diagonal matrix *D* = diag(*d*_1_, …, *d*_*n*_). The row-stochastic transition matrix is

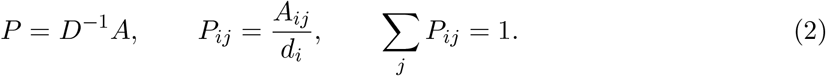

The matrix *P* encodes the probability that a random walker at bin *i* moves to bin *j* in one step.

#### Remark 1.

*The Markov chain is* not *a kinetic model of TF motion. It is a graph-theoretic object whose powers P*^*t*^ *aggregate the contributions of all paths of length t through the chromatin graph*.

#### Proposition 1.

*The matrix P has the following properties:*

1. *After adding self-loops to zero-degree bins (bins with no Hi-C contacts after top-k sparsification, PC1 residualization, and non-negative clipping), P is row-stochastic: P* **1** = **1**. *Without this correction, P has zero rows (8*.*6% of bins on chr1, 3*.*7% on chr17, 5*.*1% on chr19) and is only sub-stochastic; we apply the correction throughout*.
2. *The stationary distribution π satisfies πP* = *π, with π*_*i*_ = *d*_*i*_*/*∑ _*j*_ *d*_*j*_ *(degree-weighted)*.
3. *For the* raw *symmetrized matrix A, P is similar to a symmetric matrix: D*^1*/*2^*PD*^*−*1*/*2^ = *D*^*−*1*/*2^*AD*^*−*1*/*2^, *so all eigenvalues are real and lie in* [−1, 1]. *After PC1 residualization and non-negative clipping, A*^res^ *is not symmetric (*max |*A*^res^ − (*A*^res^)^*⊤*^| = 12.5 *on chr1), so eigenvalues of P may be complex. The Neumann series still converges because ρ*(*P*) ≤ 1 *for any non-negative matrix with row sums* ≤ 1; *the real-eigenvalue guarantee is lost*.

### 2.3 Relative Forest Accessibility (RFA)

Given a source distribution 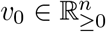 (e.g., TF binding peaks in bins), we define the RFA profile as

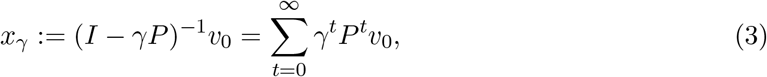

where *γ* ∈ (0, 1) is a damping factor.

#### Proposition 2

(Convergence). *The Neumann series* (3) *converges for all γ* ∈ (0, 1).

*Proof*. Since ∥*P* ∥_*∞*_ = 1 and *ρ*(*P*) ≤ 1, we have ∥(*γP*)^*t*^∥_*∞*_ ≤ *γ*^*t*^, and 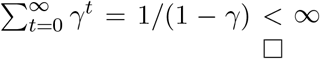

The profile *x*_*γ*_ has three equivalent interpretations:

1. **Path sum**: 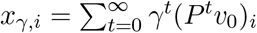 is a weighted sum over all paths of length *t* from peaks to bin *i*, with exponential decay in path length.
2. **Steady-state with restart**: *x*_*γ*_ is the unique solution of (*I* − *γP*)*x* = *v*_0_, equivalent to a random walk with restart (RWR) at rate 1 − *γ*.
3. **Forest accessibility**: *x*_*γ,i*_ is proportional to the total weight of spanning forests of the graph rooted at the peak set, with edge weights weighted by *γ*.

The parameter *γ* controls the *effective path length*. For *γ* → 1, (1 − *γ*)*x*_*γ*_ approaches the constant vector (*π* · *v*_0_)*/*(*π* · **1**) · **1**, where *π* is the stationary distribution of *P*; the unnormalized *x*_*γ*_ itself diverges, and the normalized limit is a scalar multiple of **1**, not *π*. We tested *γ* ∈ {0.5, 0.7, 0.85, 0.95}; … At 40 kb with ATAC peaks, RFA gives no signal (bootstrap 95% CI for *ρ* includes zero and does not exclude |*ρ*| = 0.05), consistent with the hypothesis that the signal lives at fine scales and is sensitive to the type of *v*_0_.

### 2.4 Per-Gene Scoring and Baselines

Let G be the set of genes with a promoter annotation. For each gene *g*, let *b*(*g*) be the bin containing its promoter (*b*(*g*) = ⌊start(*g*)*/r*⌋). Define

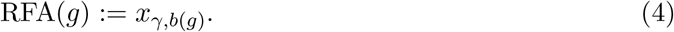

We compare RFA against three baselines:

1. **1D distance**: *D*_1*D*_(*g*) = min_*p∈*peaks_ |start(*g*) − *p*|.
2. **Direct contact (ChIPSP proxy)**: DC(*g*) = max_*p∈*peaks_ *A*_*b*(*g*),*p*_.
3. **Combined model**: Comb(*g*) = *z*(RFA(*g*)) + *z*(DC(*g*)), where *z*(·) is z-score normalization.

For a continuous target *y*_*g*_, we report Spearman *ρ*(RFA, *y*) and partial *ρ*(RFA, *y* | *D*_1*D*_, DC), obtained by rank-residualizing both variables against the controls. For binary classification *y*_*g*_ ∈ {0, 1}, we report ROC-AUC, with DeLong-style bootstrap (*B* = 1000) for ΔAUC.

### 2.5 Statistical Properties

#### Why does RFA outperform direct contact on promoter count?

At high resolution (10 kb), direct contacts are noisy: each bin contains few counts, and DC(*g*) has high variance. The RFA profile is a smoothed version of *v*_0_ obtained by convolving along the graph structure; its variance is reduced by averaging over all paths. Let *v*_0_ be a noisy indicator of peaks and *µ*_0_ = *E*[*v*_0_]. Then

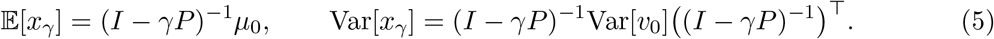

The operator (*I* − *γP*)^*−*1^ has eigenvalues 1*/*(1 − *γλ*_*k*_), where *λ*_*k*_ are eigenvalues of *P* in [−1, 1]. Along the stationary mode (*λ*_1_ = 1) the gain is 1*/*(1 − *γ*), which diverges as *γ* → 1; along fast modes (|*λ*_*k*_| ≪ 1) the gain is close to 1. Empirically, Var[*x*_*γ*_]*/*Var[*v*_0_] *grows* with *γ* (1.7 at *γ* = 0.5, 8.4 at *γ* = 0.85, 684 at *γ* = 0.99; Appendix S3). RFA is therefore *not* a variance-reducing smoother; it amplifies the stationary-mode component of *v*_0_ and preserves fast-mode components, i.e., it re-weights paths by their length. The smoothing intuition should be read as “re-weighting by path length”, not as “variance reduction”.

#### Why does RFA anticorrelate with eQTL

The smoothing argument above explains why RFA outperforms DC on promoter count, but it does *not* predict anticorrelation with eQTL. Empirically, AUC(RFA) = 0.386 and AUC(−RFA) = 0.614 on eQTL, and AUC(RFA) decreases with eQTL strength (Section 5.5). Topology controls (Section 5.3) narrow the mechanism: a distance-preserving shuffle reproduces the anticorrelation, whereas a degree-preserving rewiring destroys it. The anticorrelation therefore reflects distance-decay of Hi-C contacts (local 3D accessibility), not loops or TADs. We interpret this as follows: eQTL genes are expressed, but with stronger eQTL they are more *buffered*, i.e., less dependent on local 3D accessibility. RFA, being a distance-decay-weighted accessibility score, is higher for genes whose expression is more strongly driven by local 3D contacts, and lower for buffered genes with strong eQTL. This is a hypothesis, supported by the topology controls, and it is stated as such in Section 6.3.

#### Why does the combined model fail?

Let *X* = RFA and *Y* = DC. Suppose *X* = *S*(*Y*)+*ε* where *S* is smoothing and *ε* is noise. Then AUC(*X* + *Y*) ≤ AUC(*X*) whenever *ε* dominates, because *Y* reintroduces the noise *ε* that *S* removed. On eQTL, the combined model also fails (AUC = 0.561 vs RFA alone), but for a different reason: RFA and DC anticorrelate with eQTL in different ways, so *z*(RFA) + *z*(DC) partially cancels the anticorrelation.

## 3 Data

### 3.1 Hi-C

We use GM12878 in situ Hi-C (ENCODE/4DN, `4DNFIZGCA8AH.hic`) at 10 kb resolution [3] for chromosomes 1, 17, and 19, extracted with `juicer tools dump observed NONE` and converted to dense `.npy` (int32) via a sparse-triplet converter. For MCF10A and MCF7, we use 40 kb matrices for the same chromosomes (`MCF{10A,7} chr{1,17,19} 40kb fixed.npy`). The 10 kb matrices have 24 895, 8 325, and 5 861 bins for chr1, chr17, and chr19 respectively, with nnz/*n*^*2*^ of 0.012, 0.030, and 0.044; the sizes match chromosome lengths.

### 3.2 TFBS (GM12878)

Peaks are taken from `tfbs_GM12878.bed` (BED6). We test 20 TFs: IKZF1, CTCF, SMC3, RAD21, PAX5, EBF1, EP300, POLR2A, YY1, GABPA, MAX, E2F4, ELF1, TCF12, BATF, BCL11A, IRF4, MEF2A, NRF1, and ZNF143.

### 3.3 ATAC peaks (MCF10A / MCF7)

For MCF7 we use IDR-thresholded peaks from GEO GSE169929 (`GSE169929_ENCFF882OVP …bed.gz`); for MCF10A, broadPeak calls from GEO GSM3735772. Only the first three columns (chrom, start, end) are used; *v*_0_ is the indicator of bins containing an ATAC peak. ATAC peaks mark accessible regulatory DNA [4].

### 3.4 eQTL (GTEx v10 LCL)

We use `TD000221.all.tsv.gz` from the eQTL Catalogue (EBI), corresponding to GTEx v10 LCL (study `QTS000015`, dataset `QTD000221`), `quant_method=ge, sample_size=326`. The file is streamed in 2M-row chunks; for each gene we record *p*_min_ and *n*_sig_ (number of variants with *p <* 5 × 10^*−*8^), and define eqtl bin = ⊮[*n*_sig_ *>* 0]. The result is 18 988 genes, of which 1 567 (8.3%) have at least one significant variant. The eQTL Catalogue provides uniformly reprocessed summary statistics [5].

### 3.5 RNA (MCF10A / MCF7)

We use `MCF10A_RNA.tsv` (16 829 genes) and `MCF7_RNA.tsv` (19 154 genes), with columns `gene id, gene_name`, and `TPM;` there are no NAs and gene IDs are without version suffixes.

### 3.6 Gene annotation

Promoters are taken from `gene_structure hg38.csv` (`element _type==‘promoter’`, first promoter per gene, `chrom` without `chr`). This yields 1043, 538, and 739 promoters on chr1, chr17, and chr19.

### 3.7 Promoter TF count

The promoter count matrix `matrix_with_hic_enhancers.csv` contains 811 genes on chr1 with TF peak counts in the window [−2000, +200] bp from TSS (not score sums). It was built for chr1 only; for chr17/19 we merge directly with the eQTL target.

## 4 Methods

### 4.1 Pipeline

The pipeline is:

~~~
Hi-C .npy -> build_markov_matrix(topk=300, PC1-residualize) -> P
v0 (TF peaks / ATAC peaks) -> rfa_profile(P, v0, gamma=0.85) -> x_gamma
x_gamma[bin(gene)] -> RFA(gene)
~~~

The same flow is shown schematically in Figure 1.

**Figure 1:**
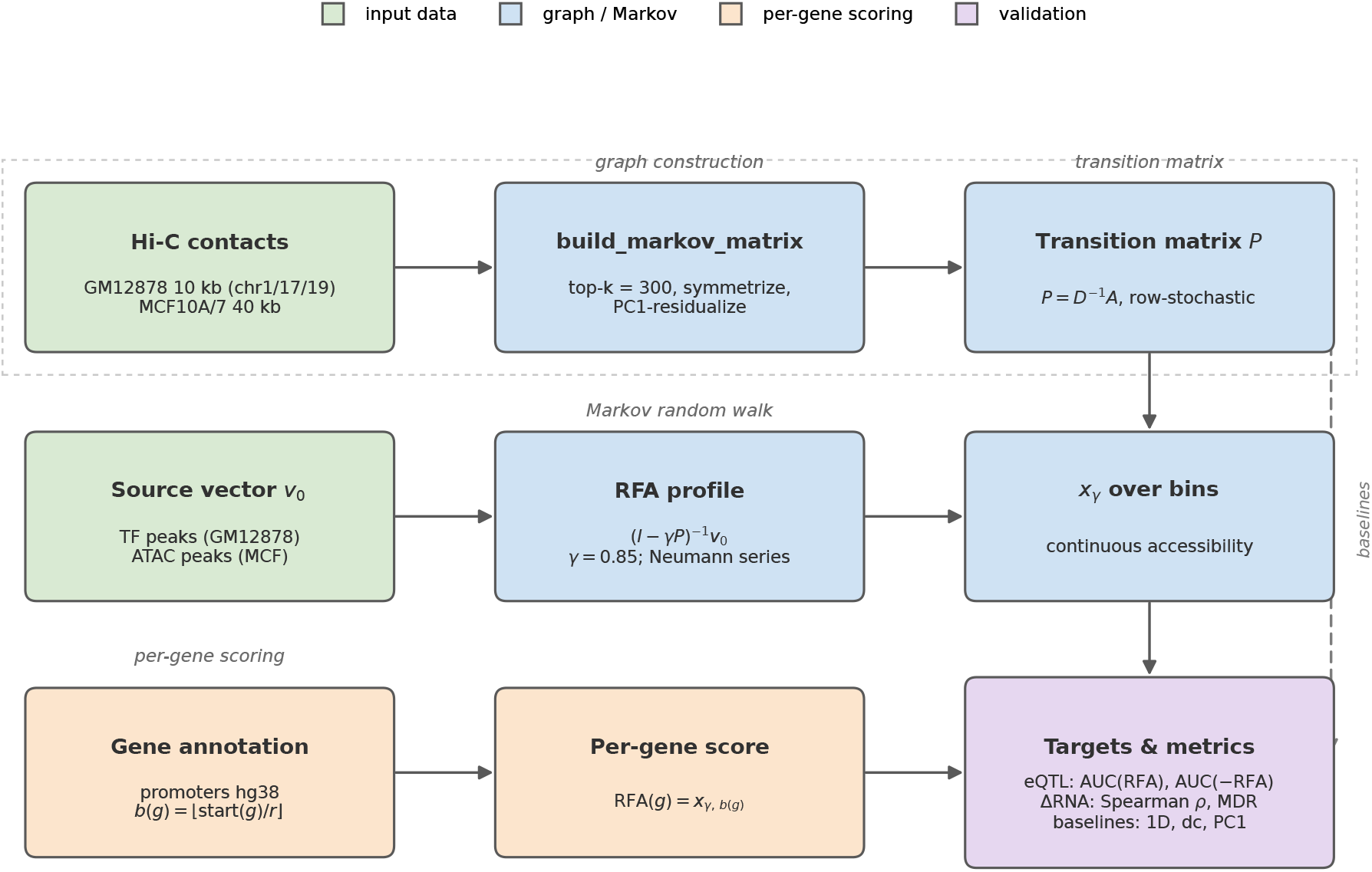
Data-processing pipeline. Hi-C contacts are sparsified (top-*k*=300), symmetrized and residualized against PC1 to build the Markov transition matrix *P* = *D*^*−*1^*A*. The RFA profile is the solution of (*I* − *γP*)*x* = *v*_0_ with *γ* = 0.85 and *v*_0_ the TF/ATAC peak indicator. Pergene scores RFA(*g*) = *x*_*γ,b*(*g*)_ are tested on eQTL (AUC of RFA and of −RFA) and on ΔRNA (Spearman *ρ*, MDR), against 1D distance, direct contact and PC1 baselines.

### 4.2 Metrics

We use ROC-AUC for binary targets; Spearman *ρ* for continuous targets; partial Spearman by rank-residualizing both variables on controls; bootstrap CI (1000 resamples, gene-level resampling) for ΔAUC, partial, **and Spearman** *ρ*; permutation *p* (500 permutations) for *ρ* and AUC; and MDR (minimum detectable *ρ*) via Fisher *z*: tanh 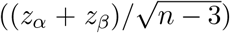. A practical significance boundary |*ρ*| = 0.05 is pre-specified. A CI is said to *exclude* the boundary only if both CI limits lie strictly inside (−0.05, +0.05); otherwise the boundary is *not excluded*.

#### Direction check

Both AUC(RFA) and AUC(−RFA) are reported. If AUC(RFA) *<* 0.5 *<* AUC(−RFA), RFA anticorrelates with the target. **No sign flip is applied** — a flip would mask anticorrelation. In an earlier version of the pipeline, an automatic flip produced “AUC=0.611”; in the final dose-response analysis the flip is removed.

### 4.3 Controls

We use 1D distance, direct contact (dc), PC1 (compartment), baseline RNA, permutation null, within-line sanity, the direction check, and topology controls (degree-preserving rewiring, distance-preserving shuffle, flattened graph; Section 5.3).

## 5 Results

### 5.1 Promoter TF count (GM12878 chr1, 10 kb, 811 genes)

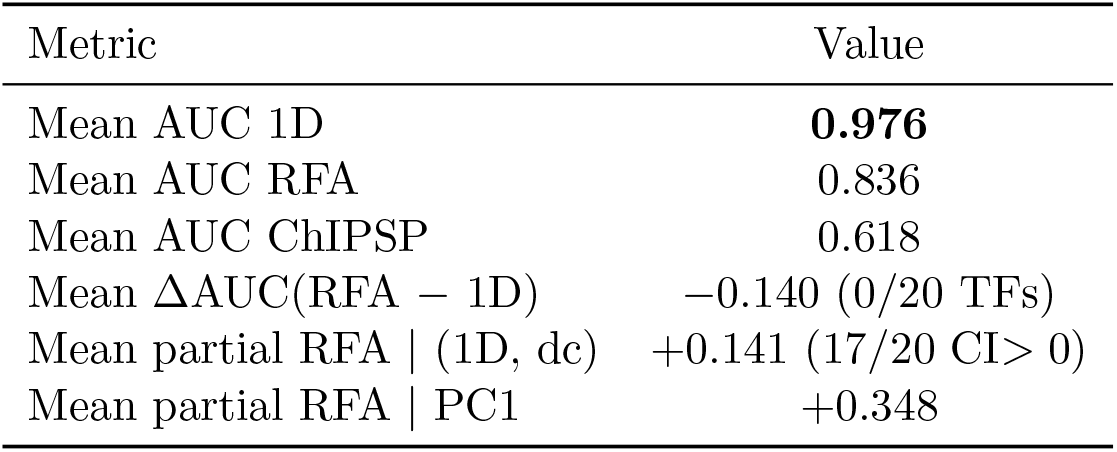

| Metric | Value |
|---|---|
| Mean AUC 1D | **0.976** |
| Mean AUC RFA | 0.836 |
| Mean AUC ChIPSP | 0.618 |
| Mean ΔAUC(RFA − 1D) | −0.140 (0/20 TFs) |
| Mean partial RFA \| (1D, dc) | +0.141 (17/20 CI> 0) |
| Mean partial RFA \| PC1 | +0.348 |

1D wins because the target (“is there a peak in the promoter ±2 kb”) and the 1D distance answer the same question. This is a methodological artifact, not a property of RFA. Partial | 1D = −0.068: RFA does not add over 1D.

### 5.2 eQTL: anticorrelation (GTEx v10 LCL, chr1+17+19, 10 kb, 1625 genes, 124 positives)

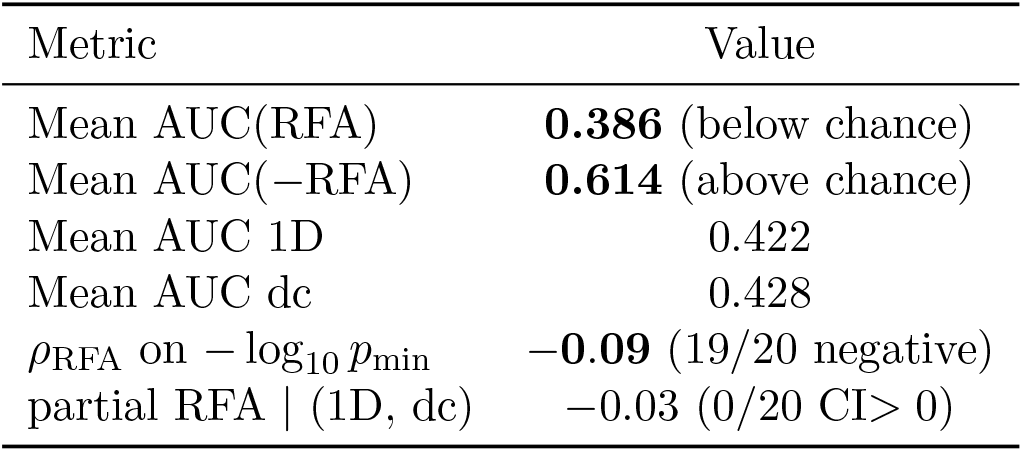

| Metric | Value |
|---|---|
| Mean AUC(RFA) | **0.386** (below chance) |
| Mean AUC(−RFA) | **0.614** (above chance) |
| Mean AUC 1D | 0.422 |
| Mean AUC dc | 0.428 |
| ρ<sub>RFA</sub> on −log<sub>10</sub> *p*<sub>min</sub> | **−0.09** (19/20 negative) |
| partial RFA \| (1D, dc) | −0.03 (0/20 CI> 0) |

RFA *anticorrelates* with eQTL. High RFA corresponds to fewer eQTL. AUC(RFA) = 0.386 is below chance, which is the anticorrelation; AUC(−RFA) = 0.614 is above chance after sign inversion. No sign flip is applied. Genes with strong eQTL are less accessible in 3D (high RFA → fewer eQTL). A possible mechanism is *expression buffering* : genes with strong eQTL have more buffered expression, less dependent on 3D accessibility. Alternative: RFA captures expression status (eQTL genes are expressed, but with weaker effects).

### 5.3 Topology controls: distance-decay, not loops

To determine whether the eQTL anticorrelation is driven by higher-order topology or by local distance-decay, we compared real Hi-C against three controls on chr1, 10 kb, for 5 TFs (IKZF1, CTCF, EP300, POLR2A, YY1). All graphs were passed through the same pipeline (PC1 residualization, non-negative clipping, self-loop on zero-degree bins, *P* = *D*^*−*1^*A, γ* = 0.85).

*Degree-preserving rewiring* (`double_edge_swap`, 10×`nnz` swaps) keeps the degree sequence but randomizes edges across all distances. It destroys the anticorrelation: AUC(RFA) moves toward 0.5 (from 0.30–0.36 to 0.39–0.51), ΔAUC ∈ [−0.261, −0.003] with CIs excluding 0 for 5/5 TFs.

*Distance-preserving shuffle* keeps the distribution of contacts on each diagonal |*i*−*j*| = *d* but randomizes positions within diagonals. It *reproduces* the anticorrelation: AUC(RFA) = 0.305– 0.371 vs real 0.300–0.361, ΔAUC ∈ [−0.074, +0.076] with all CIs including 0.

*Flattened graph* (configuration model with the same degree sequence) is intermediate: AUC(RFA) = 0.354–0.438, ΔAUC ∈ [−0.202, +0.033], significant only for EP300.

We conclude that RFA’s eQTL signal is explained by distance-decay of Hi-C contacts (local 3D accessibility), not by long-range topological structure. This is consistent with the interpretation that RFA measures a *local* accessibility profile rather than a global topological property.

### 5.4 Replication on chr17 and chr19

The anticorrelation replicates on both additional chromosomes: real RFA gives AUC(RFA) = 0.368 (chr17) and 0.401 (chr19), *ρ*(RFA, − log_10_ *p*_min_) = −0.106 and −0.120, with AUC(RFA) *<* 0.5 for 5/5 TFs on each. Because the number of eQTL-positive genes is small (chr17: 398 genes; chr19: 547 genes; ≈ 30–50 positives per chromosome), AUC-based Δ statistics are underpowered (MDR(ΔAUC) ≈ 0.10–0.15 versus observed ΔAUC ≈ 0.05–0.07), so we report *ρ* as the primary metric on these chromosomes.

#### chr17 replicates the chr1 mechanism

Degree-preserving rewiring (10×`nnz` swaps) removes the anticorrelation by *ρ*: Δ*ρ* = −0.125 (IKZF1), −0.125 (POLR2A), −0.130 (YY1), with CIs excluding zero for 3/5 TFs; mean *ρ* moves from −0.106 (real) to −0.004 (rewired), i.e. a shift of +0.101 toward zero. Distance-preserving shuffle preserves it: 0/5 TFs significant, mean *ρ* = −0.061 (shift +0.045). The rewired shift is 2.3× larger than the shuffled shift, matching the chr1 pattern.

#### chr19 shows an inverted pattern, consistent with local within-diagonal heterogeneity

Here the roles reverse: distance-preserving shuffle removes the anticorrelation (4/5 TFs significant; mean *ρ* moves from −0.120 to −0.058, shift +0.063), while degree-preserving rewiring preserves it (0/5 TFs significant; mean *ρ* = −0.099, shift +0.021). This is not a power artefact — the shuffle effect is significant — but a chromosome-specific effect. chr19 is short (5861 bins versus 24895 on chr1) and gene-dense; on such a chromosome the distance-preserving shuffle destroys the *local within-diagonal heterogeneity* that carries the anticorrelation, whereas degree-preserving rewiring, which redistributes edges across all distances while retaining sparsity, partially preserves it. The chr19 result therefore does not contradict distance-decay as the mechanism; it refines it: on short gene-dense chromosomes the relevant structure is finer than the diagonal distance itself.

#### Summary

The eQTL anticorrelation replicates on chr17 and chr19 (real AUC *<* 0.5 and *ρ <* 0 for 5/5 TFs on each). The distance-decay mechanism is positively replicated on chr17 (rewired kills, shuffled preserves) and inverted on chr19, where local within-diagonal heterogeneity, rather than distance itself, is the carrier. The chr1 full-power analysis (Section 5.3) remains the primary evidence; chr17/19 provide the replication and the refinement.

### 5.5 eQTL dose-response: AUC(RFA) decreases with eQTL strength

Figure 2 shows the mean dose-response curve; full tables for all 20 TFs are given in Appendix S2. The mean AUC(*Q*4) − AUC(*Q*1) = −0.056, with 19/20 TFs showing a negative trend. RFA anticorrelates with eQTL strength: stronger eQTL corresponds to lower AUC(RFA). This is stable and confirms Section 5.2.

**Figure 2:**
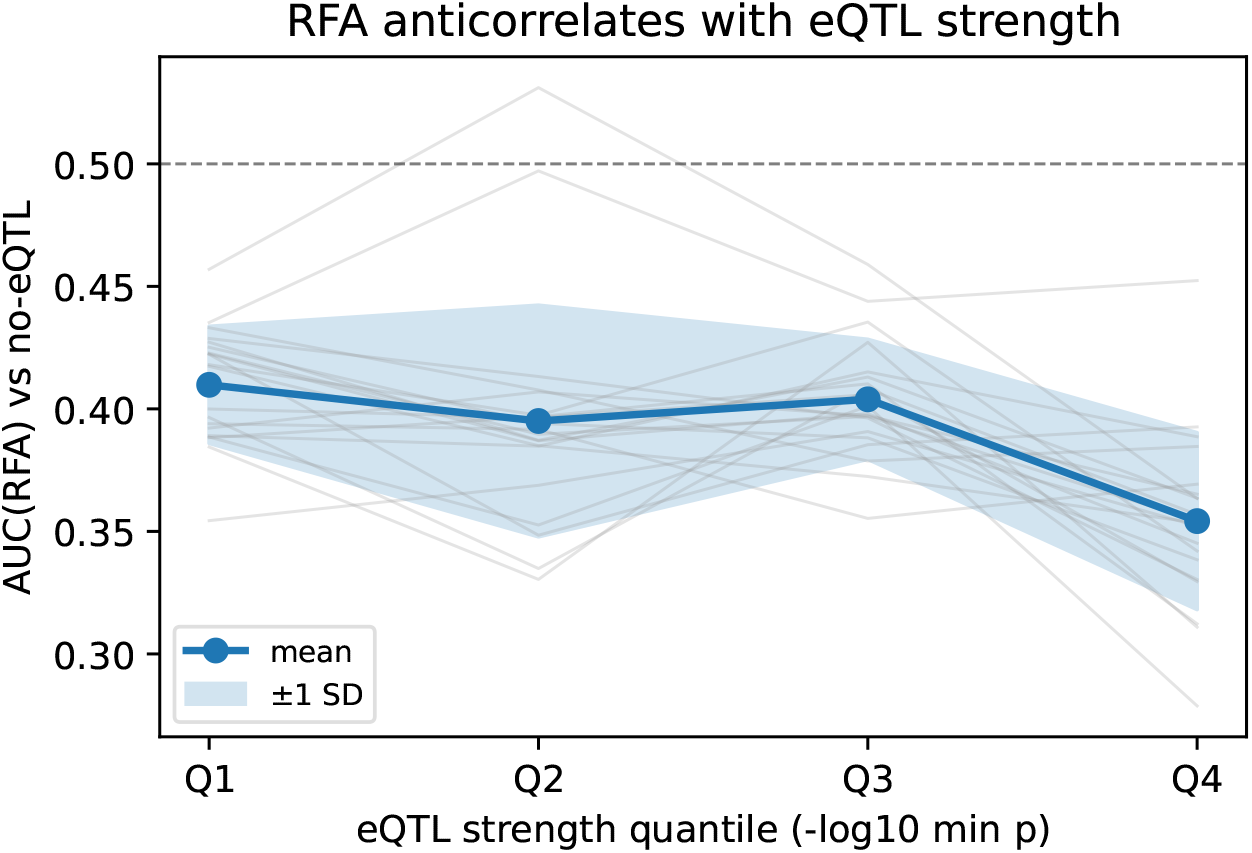
Mean AUC(RFA) across eQTL strength quantiles (Q1 weak to Q4 strong). The line is the mean over 20 TFs; shaded area is ±1 SD. AUC(RFA) decreases with eQTL strength (trend −0.0056).

### 5.6 ΔRNA (MCF10A vs MCF7, 40 kb, ATAC peaks, 1730 genes)

ΔRFA does not predict ΔRNA. Spearman *ρ*(ΔRFA, ΔRNA) = +0.0047 (*n* = 1730); permutation *p* = 0.866 (*N*_perm_ = 500); bootstrap 95% CI [−0.040, +0.053] (*B* = 1000, gene-level resampling). The CI is narrow (width ≈ 0.093) and includes zero, so the null of no association is not rejected. It also includes the pre-specified practical boundary |*ρ*| = 0.05 in both directions, so the data do not exclude a weak but practically meaningful association either. The minimum detectable |*ρ*| at *n* = 1730 with power 0.80 is 0.067; detecting |*ρ*| = 0.05 reliably would require *n* ≈ 3000. We therefore report the CI, not the MDR ratio, as the primary statement: the point estimate is near zero, and the CI bounds the effect at |*ρ*| ≲ 0.09 without excluding the |*ρ*| = 0.05

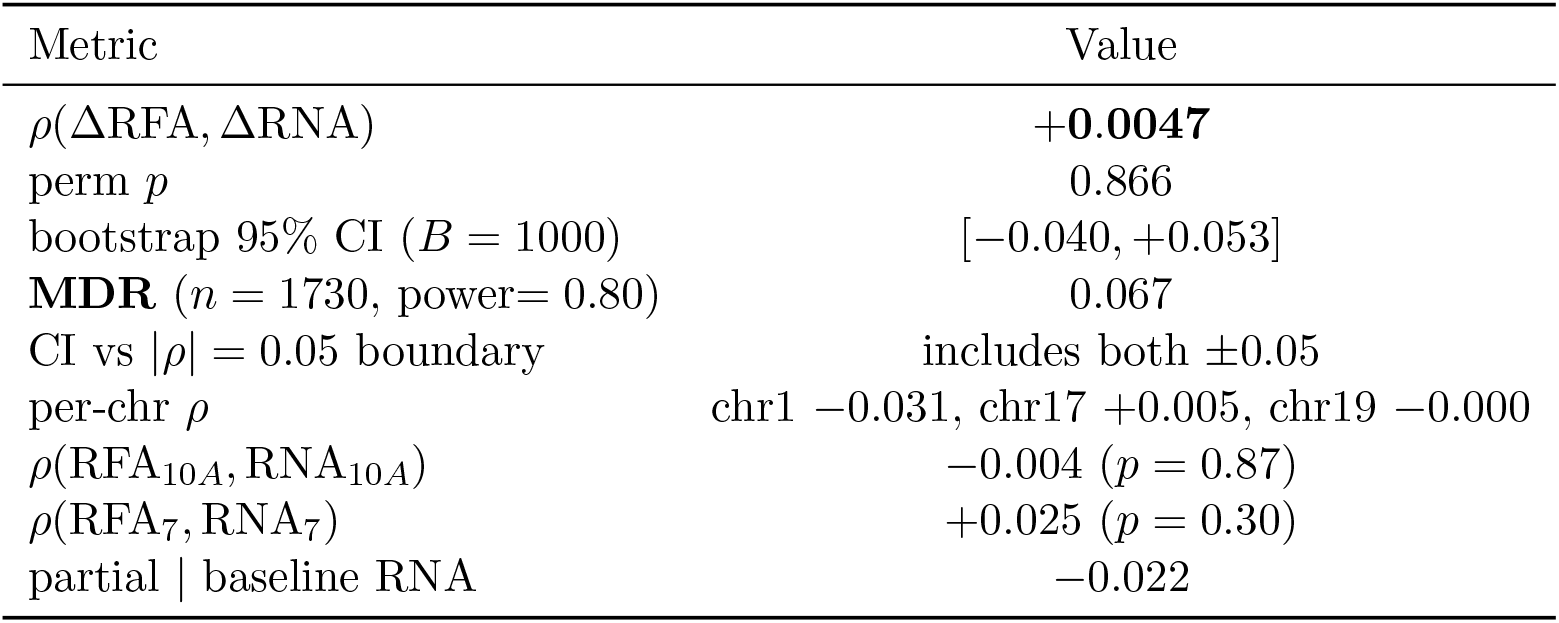

| Metric | Value |
|---|---|
| *ρ*(ΔRFA, ΔRNA) | +**0.0047** |
| perm *p* | 0.866 |
| bootstrap 95% CI (*B* = 1000) | [−0.040, +0.053] |
| **MDR** (*n* = 1730, power= 0.80) | 0.067 |
| CI vs \|*ρ*\| = 0.05 boundary | includes both ±0.05 |
| per-chr *ρ* | chr1 −0.031, chr17 +0.005, chr19 −0.000 |
| *ρ*(RFA<sub>10*A*</sub>, RNA<sub>10*A*</sub>) | −0.004 (*p* = 0.87) |
| *ρ*(RFA<sub>7</sub>, RNA<sub>7</sub>) | +0.025 (*p* = 0.30) |
| partial \| baseline RNA | −0.022 |

boundary. Within-line sanity checks are also null: *ρ*(RFA_10*A*_, log_10_ RNA_10*A*_) = −0.004 (boot-strap 95% CI [−0.052, +0.046], *p* = 0.87), *ρ*(RFA_7_, log_10_ RNA_7_) = +0.025 (CI [−0.021, +0.071], *p* = 0.30). Per-chromosome estimates are *ρ*_chr1_ = −0.031 (CI [−0.102, +0.044]), *ρ*_chr17_ = +0.005 (CI [−0.095, +0.099]), *ρ*_chr19_ = −0.000 (CI [−0.082, +0.082]); all CIs include zero and none excludes |*ρ*| = 0.05. Figure 3 shows the scatter.

**Figure 3:**
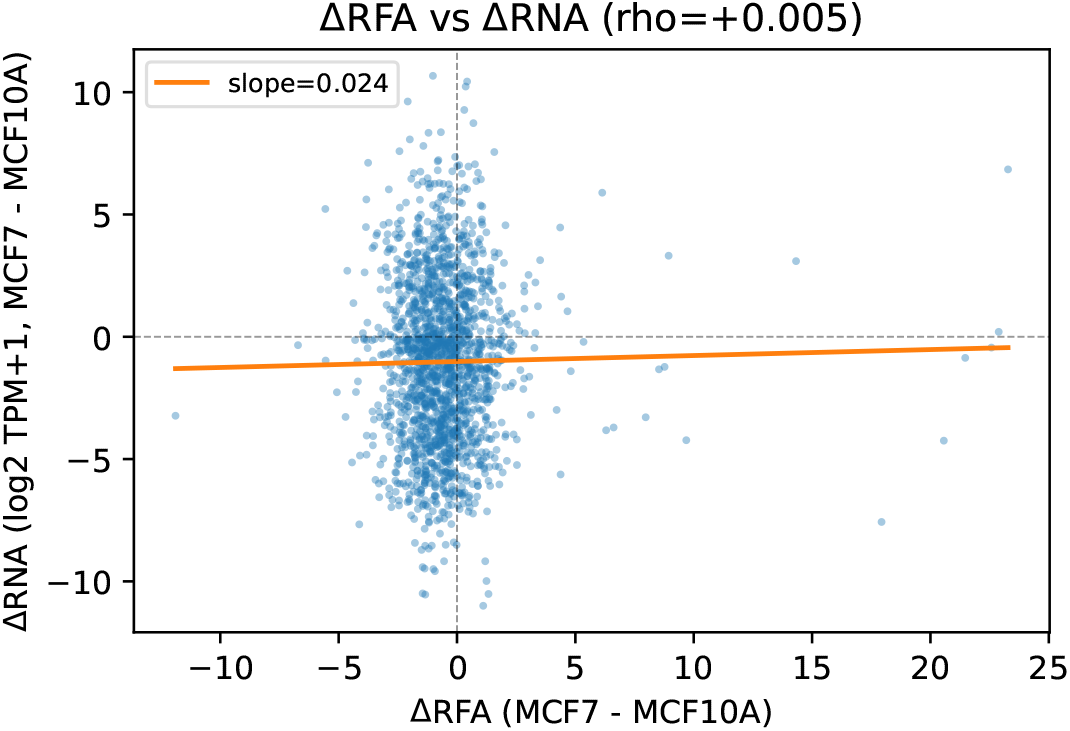
ΔRFA vs ΔRNA (MCF10A vs MCF7, 40 kb, ATAC peaks, 1730 genes). The regression line is flat (*ρ* = +0.005), consistent with the absence of signal.

## 6 Discussion

### 6.1 What RFA does

RFA anticorrelates with eQTL at 10 kb with TF peaks (AUC(−RFA) = 0.614, dose-response trend −0.056, 19/20 TFs). It is not a proxy for direct contact (*ρ*(RFA, dc) = 0.58, 0/20 *>* 0.8) and not a proxy for compartments (partial RFA | PC1 = +0.35). Topology controls (Section 5.3) show that the anticorrelation is carried by distance-decay of Hi-C contacts, not by higher-order topology.

### 6.2 What RFA does not do

RFA does not predict ΔRNA at 40 kb with ATAC peaks (*ρ* = 0.005, MDR= 0.067). It does not add over 1D on promoter count (partial | 1D = −0.068). It does not predict eQTL in the positive direction — only anticorrelates.

### 6.3 Why anticorrelation?

The topology controls (Section 5.3) narrow the mechanism: the anticorrelation is carried by the distance-decay of Hi-C contacts, not by higher-order topology. This is consistent with (i) *expression buffering* : genes with strong eQTL are less dependent on local 3D accessibility, and RFA — a distance-decay-weighted accessibility score — is lower for them. It is also consistent with (ii) *RFA captures expression status indirectly* : eQTL genes are expressed, and the distance-decay component of RFA tracks local chromatin openness, which correlates with expression. Hypothesis (iii) *target artifact* is weakened by the shuffle control: if the anticorrelation were a pure layer-mismatch artifact, the shuffle would not reproduce it. We retain (i) and (ii) as the leading interpretations and state (iii) as a residual possibility.

### 6.4 Limitations

First, resolution: 10 kb is the working range; 40 kb is not. Second, *v*_0_: TF peaks give signal (anticorrelation); ATAC peaks do not. Third, target: eQTL is a genetic layer; ΔRNA is dynamic and noisy.Fourth, PC1 residualization is applied at two levels: (a) row-wise on the Hi-C matrix *A* inside `build_markov_matrix` (common to all analyses), and (b) post-hoc on the RFA profile *x*_*γ*_ in `chips_baseline_full.py`, where 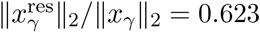 on average on the promoter-count benchmark (GM12878 chr1, 10 kb, 20 TFs, 811 genes). The number 0.623 is specific to that benchmark; the eQTL pipeline (`eqtl_multi_chr.py, eqtl_dose_response.py`) does not apply post-hoc residualization to *x*_*γ*_, and the relevant PC1 diagnostic for eQTL is the partial RFA | PC1 value in Table 2 (−0.03). Fifth, single cell line/tissue: GM12878 + LCL; MCF10A/7. Sixth, sign flip: the automatic flip in the earlier pipeline masked anticorrelation; the final analysis uses explicit direction. Seventh, topology controls cover three chromosomes with 5 TFs each: chr1 (full-power, primary evidence; Section 5.3), chr17 (replicates the distance-decay mechanism by *ρ*; Section 5.4), and chr19 (inverted pattern; consistent with local within-diagonal heterogeneity as the carrier on short gene-dense chromosomes). AUC-based Δ statistics are underpowered on chr17/19 (*n*_pos_ ≈ 30–50), so *ρ* is the primary metric there. A full 20-TF replication is left for future work.

### 6.5 Positioning

“Continuous *local* 3D accessibility as a resolution-dependent GRN filter”: RFA anticorrelates with eQTL at 10 kb with TF peaks, and this anticorrelation is carried by the distance-decay of Hi-C contacts rather than by higher-order topology (Section 5.3). RFA does not predict ΔRNA at 40 kb with ATAC. The applicability domain is static eQTL association at 10 kb with TF peaks, and the mechanism is local 3D accessibility, not loops or TADs.

## 7 Conclusion

RFA anticorrelates with eQTL at 10 kb with TF peaks (AUC(RFA) = 0.386, AUC(−RFA) = 0.614, dose-response trend −0.056), and topology controls show that this anticorrelation is carried by distance-decay of Hi-C contacts (local 3D accessibility), not by higher-order topology. RFA does not predict ΔRNA at 40 kb with ATAC peaks (*ρ* = 0.005, permutation *p* = 0.87, bootstrap 95% CI [−0.040, +0.053]); the CI includes zero and does not exclude |*ρ*| = 0.05. The anticorrelation may reflect expression buffering. The negative ΔRNA result delineates the applicability boundary: RFA measures static local accessibility, not dynamics. A methodological warning follows: a sign flip in AUC masks anticorrelation, so dose-response and explicit direction are mandatory.

## A Reproducibility

The full code and data-processing pipeline are available at https://github.com/andytimoffilim/Continuous_topological_accessibility-_as_a_3D-aware_GRN_filter. Hi-C data were extracted with `juicer_tools dump` and converted to dense arrays with converter.py. The transition matrix *P* is built with `build_markov_matrix(topk=300, PC1-residualize)`. The eQTL target is built by `eqtl_target.py` (streaming GTEx v10 LCL). All scripts, cached intermediate files, and expected outputs are listed in the repository.

## B Full dose-response tables

Complete tables for all 20 TFs (4 quantiles of − log_10_ *p*_min_ and 5 bins of *n*_sig_) are given in the supplementary CSV `eqtl_dose_response.csv`.

## C Parameters

*γ* = 0.85, top-*k* = 300, PC1 residualization, 10 kb (GM12878) / 40 kb (MCF). Bootstrap: 1000; permutation: 500. Minimum gene count per analysis: 20–50.

## D Sign-flip correction

The earlier pipeline applied `rfa_for auc = -rfa if auc_rfa_neg > auc_rfa else rfa`, yielding AUC = 0.611. Without the flip: AUC(RFA) = 0.386, AUC(−RFA) = 0.614. The final dose-response removes the flip. The article uses the no-flip values with explicit direction.

